# Spatial multi-omics enables single-cell transcriptome–metabolome inference

**DOI:** 10.64898/2026.08.06.743252

**Authors:** Lu-Yu Yang, Jing Xu, Tian-Lun Wang, Qi-Ning Zhu, Xiao-Yun Zhang, Lun-Xiu Qin, Hua-Jie Zong, Hu-Liang Jia, Xiao-Tian Shen

**Affiliations:** Department of General Surgery, Huashan Hospital, Fudan University, 12 Urumqi Road (M), Shanghai 200040, China; Cancer Metastasis Institute, Fudan University, Shanghai, China; Microsoft Co Seattle, USA; Department of Dermatology, Renji Hospital, School of Medicine Shanghai JiaoTong University 160 Pujian Road, Shanghai 200127, China

**Author notes:** To whom correspondence should be addressed: Xiao-Tian Shen, Department of General Surgery, Huashan Hospital, Cancer Metastasis Institute, Fudan University, 12 Urumqi Road (M), Shanghai 200040, China.; Hu-Liang Jia, Department of General Surgery, Huashan Hospital & Cancer Metastasis Institute, Fudan University, 12 Urumqi Road (M), Shanghai 200040, China., Hua-Jie Zong,Department of General Surgery, Huashan Hospital, Fudan University, 12 Urumqi Road (M), Shanghai 200040, China. These authors contributed equally to this work (Lu-Yu Yang, Jing Xu, Tian-Lun Wang). **Author Contributions:** Xiao-Tian Shen, Hu-Liang Jia and Hua-Jie Zong designed and supervised the study; Lu-Yu Yang, Qi-Ning Zhu and Tian-Lun Wang performed the experiments, Xiao-Tian Shen, Jing-Xu and Xiao-Yun Zhang write the code and provided helps in experimental techniques as well as data analysis, Qi-Ning Zhu and Lun-Xiu Qin helped revise the manuscript, Lu-Yu Yang prepared the manuscript.

**Keywords:** single-cell metabolomics, spatial multi-omics, multiple instance learning, transformer, MALDI-MSI, hepatocellular carcinoma, macrophage polarization

## Abstract

Joint single-cell transcriptomic–metabolomic profiling remains technically intractable. Here we present CHIMERA (**C**ell-level **H**ybrid **I**nference of **M**etabolome **E**mbedded on **R**NA **A**tlas), a data-driven framework that learns transcriptome-to-metabolome mappings from spatially paired multi-omics data and transfers them to unpaired scRNA-seq. CHIMERA generates quantitative, database-independent single-cell metabolite abundances and, by pairing them with the measured transcriptome of the same cells, enables joint co-embedding of genes and metabolites for the discovery of differential metabolites and co-regulated gene–metabolite modules. Using 10x Visium paired with MALDI-MSI from murine liver sections and a matched scRNA-seq reference, CHIMERA achieves a per-metabolite median Pearson r = 0.285 with positive cross-section generalization. On an independent Liver Cell Atlas Western-diet cohort, CHIMERA recovers metabolic reprogramming that recapitulate published non-alcoholic fatty liver disease pathophysiology. Applied to a Rarres2 (chemerin) knock-down hepatocellular carcinoma model, CHIMERA uncovers metabolic heterogeneity among tumour-associated macrophages, resolving four metabolic subclusters (MC-0 to MC-3); Rarres2 appears to drive macrophage polarization from an LAM-like MC-3 state toward Spp1+ like MC-0/MC-2 by modulating a co-regulated gene–metabolite module—a dual-omics phenotype undetectable by either modality alone. CHIMERA is the first data-driven framework for quantitative single-cell metabolome inference, opening joint transcriptomic– metabolomic analyses inaccessible to either experimental or knowledge-based computational approaches.

## Introduction

The integration of multi-omics data has become a central strategy in modern biology, enabling the discovery of biological phenomena that are invisible to any single molecular layer alone[1] [2]. By jointly analysing genomic, transcriptomic, proteomic and metabolomic information, researchers can construct comprehensive molecular portraits of cellular states and intercellular interactions[3]. Among these modalities, single-cell RNA sequencing (scRNA-seq) has matured into a routine technology, with large-scale atlases now encompassing millions of cells across diverse tissues, species and disease contexts[4-6]. Metabolomics, which captures the functional endpoints of cellular biochemistry, provides the most proximate readout of phenotypic state[7] . However, simultaneously profiling the transcriptome and metabolome of the same single cell remains a formidable technical challenge.[8, 9] . A recent proof-of-concept method, Metcell, demonstrated simultaneous metabolome and transcriptome profiling from individual liver cells, but its reliance on manual nano-capillary sampling limits scalability [10, 11]. Consequently, joint single-cell transcriptomic–metabolomic datasets remain scarce and small in scale, constituting a major bottleneck for understanding metabolic heterogeneity at cellular resolution.

Recent advances in deep learning and transformer-based models have demonstrated that information-rich molecular modalities can be computationally predicted from more accessible measurements, effectively performing cross-modal “translation”. In computational pathology, deep-learning frameworks such as HEX and HistoPlexer have shown that spatially resolved protein expression profiles encompassing dozens of biomarkers can be accurately imputed from standard haematoxylin and eosin (H&E) histology images [12, 13] . Similarly, methods including ST-Net, HisToGene, THItoGene and GHIST predict spatial gene expression from histological images by learning morphology–transcriptome associations [14-16]. In epigenomics, frameworks such as EPCOT have demonstrated that chromatin organization, histone modifications and transcription factor binding can be predicted across cell types from chromatin accessibility data alone [17]. These successes collectively establish a paradigm in which paired multi-modal data are used to train models that subsequently generalize to samples where only one modality is available.

Several computational approaches have been developed to infer metabolic states from scRNA-seq data, but they operate under fundamentally different assumptions and produce qualitatively different outputs than direct metabolite prediction. Constraint-based methods, exemplified by Compass [18], construct cell-type-specific genome-scale metabolic models and estimate reaction fluxes by optimizing flow through the metabolic network subject to gene expression constraints. scFEA employs a graph neural network on a reconstructed metabolic factor graph, modelling the non-linear relationship between enzymatic gene expression and reaction rates to infer cell-wise metabolic flux [19]. METAFlux extends flux-balance analysis with machine learning to characterize metabolic reprogramming in both bulk and single-cell settings [20]. Pathway-level scoring approaches such as scMetabolism and ssGSEA provide per-cell activity scores for predefined metabolic gene sets but do not yield quantitative metabolite-level predictions [21, 22]. MEBOCOST takes a distinct approach by inferring metabolite-mediated cell–cell communication from enzyme and sensor expression, rather than predicting intracellular metabolite abundances [23] . While these methods have yielded valuable biological insights, they share critical limitations: all rely heavily on curated metabolic pathway databases (e.g., KEGG, Recon), restricting predictions to known enzymatic reactions; their outputs are relative fluxes or pathway scores rather than metabolite abundance directly comparable to mass-spectrometry measurements; and crucially, none leverages experimentally measured metabolomic data for model training, meaning they cannot learn transcriptome–metabolome relationships that extend beyond pre-encoded biochemical knowledge.

The rapid development of spatial omics has created new opportunities for bridging the transcriptome–metabolome gap. Spatial transcriptomics platforms such as 10x Visium, Visium HD and Stereo-seq enable genome-wide gene-expression profiling with preserved tissue context, while MALDI-MSI provides spatially resolved metabolite detection at comparable or higher resolution. Critically, recent experimental workflows have demonstrated that both modalities can be acquired from the same or adjacent tissue sections. Vicari *et al*. established a same-section protocol combining MALDI-MSI and Visium on standard glass slides[24]. Hendriks *et al*. further advanced single-section integration by combining MALDI-MSI with Xenium spatial transcriptomics at pixel-scale resolution[25]. On the computational side, co-registration algorithms MAGPIE[26], MIIT [27] and SpatialMETA[28] align the distinct coordinate systems of spatial transcriptomics and mass-spectrometry imaging. Of particular relevance to the present work, haCCA integrates spatial transcriptomes and MALDI-MSI metabolomes by identifying highly correlated gene–metabolite feature pairs and performing modified spatial morphological alignment[29]. These advances collectively provide spatially co-registered transcriptomic–metabolomic datasets at near-cellular resolution, offering an unprecedented training substrate for learning direct mappings between gene expression and metabolite abundance. However, leveraging such paired spatial data to train models that can subsequently predict single-cell metabolomes from scRNA-seq data alone remains an open and largely unaddressed challenge.

Here we present CHIMERA (Cell-level Hybrid Inference of Metabolome Embedded on RNA Atlas), a data-driven framework that transfers metabolomic information from spatially paired transcriptome–metabolome measurements onto individual cells profiled by scRNA-seq. By casting each spatial spot as a probabilistic mixture of cell types and learning the transcriptome-to-metabolome mapping at cell-type resolution through multiple instance learning, CHIMERA generates quantitative, database-independent single-cell metabolome predictions directly comparable to mass-spectrometry measurements—without relying on curated pathway annotations. Applied to three murine liver sections (37,700 spatial spots) and an scRNA-seq reference of 87,554 cells, CHIMERA achieves a spot-level test R^2^ of 0.752 with positive cross-section generalization to a held-out replicate. External validation on an independent Liver Cell Atlas cohort recapitulates published NAFLD metabolite signatures, while mechanistic application to a Rarres2 knock-down hepatocellular carcinoma model reveals that chemerin/CMKLR1 signalling drives tumour-associated macrophages polarization to coordinated lipogenic, glycolytic and oxidative activation, illustrating how joint single-cell transcriptomic–metabolomic analysis enables biological insights inaccessible to either modality alone.

## Methods

### Overview

CHIMERA operates through six sequential stages: (i) preprocessing of haCCA-aligned paired spatial data into separate gene and metabolite feature matrices; (ii) cell-type deconvolution of each spot via cell2location; (iii) ridge-regression estimation of per-cell-type pure metabolic spectra under a linear-mixing prior; (iv) MIL training of a gene transformer encoder that maps cell-type-specific virtual pure transcriptomes to metabolic profiles, aggregated by deconvolution proportions; (v) spatially-blocked benchmarking against cell-type mean and ridge regression baselines; (vi) direct single-cell inference on scRNA-seq cells.

### Data preparation

#### Spatial multi-omics training data

The model was trained on co-registered Visium and MALDI-MSI data from adjacent sections of Padi4-knockout and wild-type ICC tumors (hereafter KO(16616 spots) and WT(9830 spots)). An unseen replicate *WT2* with both new Visium and new MALDI-MSI acquisitions adjacent to the WT (11741 spots). A WT3, profiled by new Visium against a shared MALDI-MSI reference with WT (15610 spots). After alignment each spot *S* carries a gene-expression vector ***t***_*s*_ ∈ ℝ^*G*^ sand a metabolite/*m/z* intensity vector ***m***_*s*_ ∈ ℝ^*F*^ where *G* denotes the number of genes and *F* the number of measured features.

#### scRNA-seq reference

A mouse scRNA-seq dataset from a YAP/AKT-driven hepatocarcinogenesis study comprising 87,554 cells × 31,253 genes was used as the single-cell reference. Cells were annotated into nine cell types (B/Plasma, T/NK, Mesenchyme, Mono/Macrophage, Neutrophil, DC/Mast, and three additional hepatocyte subtypes), denoted *K =* {1,…,*K* } with *K=*9.

#### External validation cohorts

Two publicly available mouse liver scRNA-seq datasets were used for external validation: (i) a Liver Cell Atlas subset of CD45-hepatic cells(n=33214) profiled under standard diet (SD) and Western diet (WD) at 24 and 36 weeks[30] (denoted NDWD cohort); and (ii) an in-house *Rarres2* (chemerin) knock-down hepatocellular carcinoma (HCC) model with 40995 cells comprising *Rarres2* knock-down (KD) and normal control (NC) animals. Neither cohort overlaps with the training data in animals, samples or measurement batch.

#### Preprocessing

Gene and metabolite features were separated from the merged haCCA-aligned matrices using nomenclature rules: metabolite names were identified by chemical notation (lipid class prefixes, fatty-acid chain annotations, systematic acid/carnitine suffixes), while remaining short alphanumeric symbols were retained as genes. Metabolite intensities were kept in raw (non-log) space; gene expression was log1p-normalized. A common gene set of 15054 genes shared across WT, KO and the scRNA-seq reference was used throughout. Of 212 total metabolite features, 189 with reliable structural annotations were retained for modelling.

#### Cell-Type Deconvolution

Cell-type composition at each spatial spot was inferred using cell2location in a two-step procedure. In step one, a negative-binomial regression (RegressionModel) was fitted to the scRNA-seq reference (raw counts, cell-type label, sample identity as batch covariate) to estimate per-cell-type average expression signatures 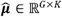. In step two, the spatial model was fitted to each tissue section separately.The prior on cells per spot was set to *N*_cells_ = 10. Output is a cell-type proportion matrix Π ∈ ℝ^*S*×*K*^, where *π*_*sk*_ is the estimated abundance of cell type *k* at spot *s*.

### Pure Spectrum Estimation

Under a linear mixing assumption the observed metabolite profile at spot *s* is modeled as

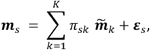

Where 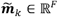 is the latent *pure spectrum* of cell type *k*. Stacking spots into matrices ***M*** ∈ ℝ^*S*×*F*^and Π ∈ ℝ^*S*×*K*^, the pure-spectra matrix 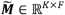 is estimated by ridge regression:

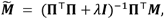

with *λ* selected by generalized cross-validation. Training was performed on the combined WT and KO sections (n=25959 spots). For the metabolite route 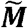 has shape 9*189.

### MIL Model Architecture

#### Gene transformer encoder

For each cell type *k* we learn a mapping *g*: ℝ^*H*^ → ℝ^*F*^ from a highly-variable-gene (HVG) expression profile to a metabolic profile, with a single shared encoder across all cell types. The primary configuration uses H = 256 HVGs (*v2*), selected by variance on the training transcriptome from the 15054 common genes. An auxiliary variant with H = 512 HVGs (*v2_512*) is retained for external-validation analyses (see below). Each gene *h* is encoded as a token that fuses

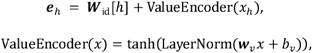

where ***W***_id_ ∈ ℝ^*H*X*D*^ (*D* = 256) is a learned gene-identity embedding and *w*_*V*_,*b*_*v*_ ∈ ℝ^*D*^ map the scalar expression to . ℝ^*D*^ A learnable [CLS] token ***c*** ∈ ℝ^*D*^is prepended; the output at position 0 after *N*_layer_ =6 pre-norm transformer encoder layers (*N*_head_ = 4, *d*_*ff*_ = 1024, GELU, dropout 0.1) serves as the cell-type metabolic embedding *z*. A two-layer MLP with LayerNorm, GELU and dropout decodes *z* to the *F*-dimensional metabolic profile. Identity embeddings and the CLS token are initialized from *N* (0,0.02); linear layers use Xavier-uniform with zero biases. The primary HVG-256 model totals -5□M parameters.

#### Virtual pure spot augmentation

To provide cell-type-resolved supervision, we construct *virtual pure spots* for each cell type *k* at spot *s*:

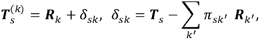

where ***R***_*k*_ is the average HVG expression of cell type *k* in the scRNA-seq reference, ***T***_S_ is the observed spot transcriptome, and *δ*_*sk*_ is a spot-specific residual that preserves local transcriptomic context while isolating cell-type identity. The virtual pure spot serves as the per-cell-type input to the encoder during training.

#### MIL prediction and loss

The predicted spot metabolome is

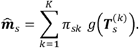

Training minimizes a mixed objective combining mean-squared error with a Pearson correlation penalty:

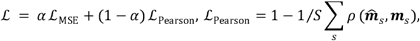

with *α* = 0.5. Optimization used AdamW with cosine-annealing learning-rate schedule, batch size 32, maximum 400 epochs and early stopping (patience 50) on the per-metabolite median validation *R*^2.^

### Single-Cell Inference

After training, the metabolome of a single cell *n* in the scRNA-seq dataset with HVG expression *z*_*n*;_ ∈ ℝ^*H*^ is predicted directly as

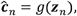

bypassing MIL aggregation since each cell represents a pure cell-type observation. Predictions for cells were exported as AnnData objects and imported into Seurat as new assays for downstream analysis.

### Spatial Cross-Validation

To prevent inflation of performance estimates due to spatial autocorrelation, spots were partitioned by pairwise spatial distance into three non-overlapping zones: training (60%), gap (10%, buffer excluded from both training and evaluation) and test (30%). A second held-out tissue section (WT2) was used as an unseen-sample generalization cohort.

### Evaluation Metrics

Let *m*_*sf*_ and 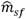 denote the observed and predicted intensity of metabolite *f* (*f* = 1,…,*F*; *F* = 189) at spot *s* (*s* = 1,…, *S*). All metrics were evaluated on the internal WT+KO test fold, the WT2 replicate and the WT3 section.

#### Coefficient of determination (R^2^)

R^2^ measures the fraction of variance explained relative to a per-feature mean baseline.

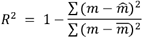

The overall spot-level R^2^ pools all spot–metabolite pairs; the per-metabolite R^2^ is computed for each metabolite across spots and summarized by its median over the *F* metabolites; the fraction *R*^2^ > 0 is the proportion of metabolites predicted better than their per-metabolite mean.

#### Pearson correlation (r)

r measures the linear agreement between predicted and observed profiles, independent of scale.

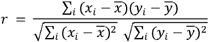

The per-spot r correlates the predicted and observed profiles across the *F* metabolites within each spot (median over spots); the per-metabolite r correlates prediction and observation across the *S* spots for each metabolite (median and IQR over metabolites, with significance Bonferroni-corrected).

### Baseline

We evaluated a ridge (L2-regularized linear) regression baseline under the same spatial cross-validation splits as CHIMERA. Gene expression was first reduced to its leading 50 principal components; a ridge regression was then fit from these components to the 189-dimensional metabolite vector by leave-one-out cross-validation.

### Ablation of Gene-Feature Selection Strategies

Eight alternative gene-feature strategies were compared under identical training and evaluation pipelines: (i) **HVG-256 (v2)**: 256 top-variance genes from the 15,054 common gene set (primary model); (ii) **HVG-512 (v2_512)**: 512 top-variance genes (deeper variant); (iii) **CorrMet-256**: 256 KEGG metabolic genes with highest Pearson correlation to measured metabolites; (iv) **CorrMet-AllGene-256**: 256 genes (any pathway) with highest metabolite-correlation magnitude;(v) **Hybrid-256**: concatenation of 128 HVG + 128 CorrMet genes; (vi) **MetGene-256**: KEGG-variance-selected 256 metabolic genes; (vii) **Bottleneck-256**: a linear 512-256 projection of HVG-512 tokens; (viii) **Flash-Linear**: a linear 15,054-to-256 projection bypassing the transformer. Each strategy was evaluated on the per-metabolite median Pearson r.

### External Validation Procedures

#### NDWD cohort (diet effect)

The model was applied to 33241 CD45-hepatic cells from the Liver Cell Atlas across four conditions (SD-24w, WD-24w, SD-36w, WD-36w), with HVG alignment via the 15,054 common gene set. Differential metabolite abundance between WD and SD was tested per cell type using Wilcoxon rank-sum followed by Benjamini–Hochberg correction (FDR<0.1, log2FC>0.5. Temporal progression was assessed by comparing 24w and 36w effect-size distributions. Functional category annotations were assigned using LIPID MAPS and KEGG metabolite classes; pathway-level over-representation was tested using gseapy Fisher exact test with BH correction against a custom background of all 189 detected metabolites mapped to KEGG metabolite sets.

#### RARRES2 KD HCC cohort

The model was applied to 40995 cells from the *Rarres2* KD HCC model. KD-*vs*-NC differential testing was performed as above per cell type. Cross-model reproducibility was evaluated by intersecting significant hits from HVG-512 with those from CorrMet-256 and MetGene-256 variants, assessing both significant-hit overlap and directional concordance.

### Macrophage Metabolic Subclustering and Trajectory

Among 40995 cells, 8505 were annotated as macrophages (Cd5l^+^ Vcam1^+^ or Spp1^+^ subtypes). The top 30 KD-*vs*-NC differentially abundant metabolites (ranked by FDR) were used to cluster macrophages. Data were scaled, reduced to 15 principal components, built into a 30-nearest-neighbour graph, and clustered by the Leiden algorithm at resolution 0.09 using scanpy. Four metabolic subclusters (MC-0 to MC-3) were recovered. Pseudotime was estimated by diffusion pseudotime (DPT) with the root cell placed in MC-3. Gene set over-representation analysis was performed per cluster using gseapy against the KEGG with a custom background of 32589 detected genes.

For the trajectory-ordered comparison, metabolites and transcriptomic pathway scores were correlated with DPT rank across the four clusters (Spearman *ρ*). KD-*vs*-NC chi-square tests on cluster composition were performed with Yates continuity correction; macrophage subtype proportion shifts were visualized using stacked bar plots.

### Statistical analysis

Differential metabolite abundance between conditions was assessed by two-sided Wilcoxon rank-sum tests with Benjamini–Hochberg correction (FDR<0.1, log2FC>0.5); Pathway enrichment was tested by one-sided Fisher exact tests with Benjamini–Hochberg correction against a custom background of 189 detected metabolites (or, for gene-level enrichment, 32,589 detected genes). Contingency comparisons of cell-cluster proportions between conditions used^2^ tests with df = 3 and Yates continuity correction. All statistical tests were performed in SciPy v1.11.1 and statsmodels v0.14.0.

### Hardware

Model training for the primary HVG-256 configuration was performed on a single NVIDIA GeForce RTX 5070 GPU (12 GB VRAM) housed in a workstation with a 16-core Intel Core i9 CPU and 64 GB DDR5 RAM running Windows 11 (Python 3.10 via Miniconda). Training the primary HVG-256 transformer to convergence (epoch 143) required approximately 4.5 GPU-hours at ∼ 110 s per epoch (batch size 32). The larger HVG-512 (v2_512) variant and all other 512-gene ablations were trained on a cloud-based NVIDIA RTX 4090 GPU (48 GB VRAM), requiring approximately 8–12 GPU-hours per route at ∼ 3–4 min per epoch. Cell-type deconvolution with cell2location was performed on a local NVIDIA RTX 4090 (24 GB VRAM) and required approximately 5 GPU-hours for 30,000 spatial training epochs. Code and trained model weights are available at https://github.com/shenxiaotianCNS/CHIMERA.

## Results

### CHIMERA: a multiple instance learning framework for single-cell metabolome prediction

Single-cell transcriptomic atlases now span millions of cells, yet no metabolomic platform approaches comparable scale. CHIMERA closes this gap by learning a transcriptome-to-metabolome mapping from spatially paired Visium and MALDI-MSI data, and transferring it to unpaired scRNA-seq (Fig. 1a).

**Figure 1:**
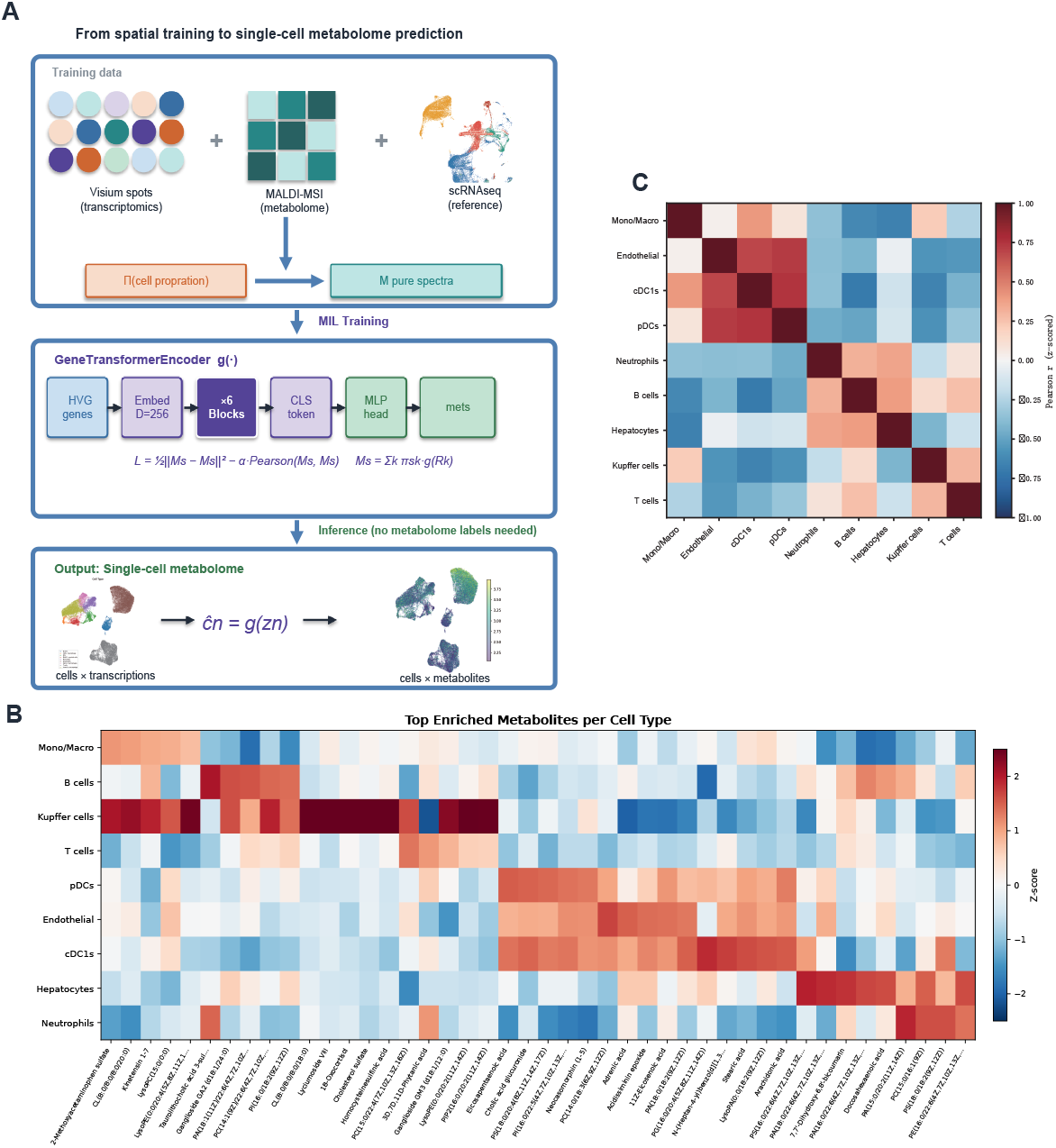
CHIMERA: an MIL-based framework for predicting single-cell metabolomes from transcriptomic input. A: Overview of the CHIMERA workflow. Spatially paired Visium transcriptomics (spot-level transcriptome *t*_*s*_ ∈ ℝ^*G*^ and MALDI-MSI metabolomics *m*_s_ ∈ ℝ^*F*^, together with a matched scRNA-seq reference providing cell-type labels k, are used to train a transformer-based mapping *g*: ℝ^*H*^ ℝ^*F*^ from gene expression to metabolite abundance. Cell-type proportions Π ∈ ℝ^*S*×*K*^ are inferred with cell2location, and a pure-spectrum matrix *M* ∈ ℝ^*k* ×*F*^ is recovered by ridge regression. The GeneTransformerEncoder tokenizes H highly variable genes, passes them through six transformer blocks, aggregates via a [CLS] token, and decodes to the F-dimensional metabolic profile through a MLP head. Training follows a multiple instance learning (MIL) formulation in which the spot-level metabolome is supervised as the Π-weighted sum of per-cell-type predictions, 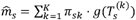, optimized by a mixed objective ℒ = *α* ℒ_*MSE*_ + (1 −*α*) ℒ_*Pearson*_. At inference, the trained encoder is applied directly to unpaired scRNA-seq transcriptomes .:*z*_*n*_ ∈ ℝ^*H*^ to yield a cell-by-metabolite matrix *ĉ*_*n*_ = *g* (*z*_*n*)_. B: Heatmap of the top enriched metabolites per cell type (z-scored across cell types), derived from the ridge-regressed pure-spectrum matrix M on the training data. C:Pairwise Pearson correlations between cell-type metabolomic profiles (z-scored).

The framework rests on three elements. First, cell-type proportions *Π* ∈ ℝ^*S*×*K*^ at each Visium spot were inferred with cell2location, and a per-cell-type pure-spectrum matrix *M* ∈ □^*K*×*F*^was recovered by ridge regression under a linear mixing assumption *m*_*s*_= Σ_*k*_ *π*_*sk*_ *m*_*k*_ + *ε*_*s*_ (Fig. 1b). Second, prediction was cast as multiple instance learning: each spot constitutes a bag of K cell-type instances, and the spot-level metabolome is supervised as the Π-weighted sum of per-cell-type predictions 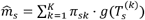, trained with a mixed MSE– Pearson objective ℒ = *α* ℒ_*MSE*_ + (1 − *α*)ℒ_*Pearon*_. This formulation extracts cell-type-resolved signal from spot-level labels without requiring pure-cell-type metabolomic standards, which are experimentally inaccessible. Third, the mapping *g*: ℝ^*H*^ → ℝ^*F*^is a six-block transformer encoder over tokenized highly variable genes, in which each gene is encoded as a learned identity embedding scaled by a magnitude-encoded expression value and aggregated into a cell-level representation via a learned [CLS] token. By decoupling gene identity from gene order, this design captures context-dependent transcriptome–metabolome relationships without pathway priors—a fundamental departure from knowledge-based metabolic inference tools. Virtual pure-spot augmentation further supplies cell-type-specific inputs while preserving local transcriptomic context.

At inference time, the trained encoder is applied directly to unpaired scRNA-seq transcriptomes *z*_*n*_ ∈ ℝ^*H*^ to yield single-cell metabolomes *z*_*n*_ ∈ ℝ^*H*^ bypassing MIL aggregation since each cell already represents a single-cell-type observation (Fig. 1a, bottom).

The model was trained on co-registered Visium and MALDI-MSI data from adjacent sections of *Padi4*-knockout and wild-type ICC tumors (hereafter KO and WT). An additional Visium section, adjacent to the WT MALDI-MSI slice, was integrated with the same WT metabolomic data to serve as an independent held-out validation set (WT2). Matched scRNA-seq was generated from tumors of matched genotype and used as the cellular reference(figure S1). Cell2location deconvolution resolved nine cell types across the WT, WT2 and KO sections; per-cell-type metabolomic profiles derived from the ridge-regressed pure-spectrum matrix revealed pronounced inter-cell-type heterogeneity (Fig. 1b), and pairwise correlation analysis showed coherent grouping of myeloid populations (Mono/Macro, cDC1s, pDCs) while Kupffer cells and hepatocytes exhibited the most distinctive metabolic signatures (Fig. 1c), reflecting their divergent metabolic roles within the hepatic tumour microenvironment.

### Systematic ablation identifies HVG-256 with MIL supervision as the best strategy that generalizes across sections

To isolate the design choices that underpin generalization, we benchmarked five gene-feature strategies under identical training and evaluation pipelines. **HVG-256** and **HVG-512** select the 256 and 512 top-variance genes from the 15,054-gene common set, respectively, and are knowledge-agnostic baselines. **CorrMet-256** selects the 256 KEGG metabolic genes with the highest Pearson correlation to measured metabolites across training spots. **MetGene-256** selects 256 KEGG metabolic genes by variance alone. **CorrMet256-AllGene** selects the 256 genes—drawn from any pathway—with the highest metabolite-correlation magnitude, thereby relaxing the KEGG constraint of CorrMet-256(Table S1). All five share identical transformer backbone, training schedule and loss.

Each strategy was evaluated on three cohorts of increasing distributional shift: an internal test fold held out from WT+KO (3,114 spots), an unseen replicate *WT2* profiled by independent Visium and MALDI-MSI acquisitions (11,741 spots), and *WT3*, profiled by new Visium against the original MALDI-MSI reference (15,610 spots). Performance was quantified by median Pearson *r* and median Spearman *ρ*, on both raw and spatially smoothed predictions (Gaussian σ = 1.5 × median nearest-neighbour distance; Fig. 2).

**Figure 2:**
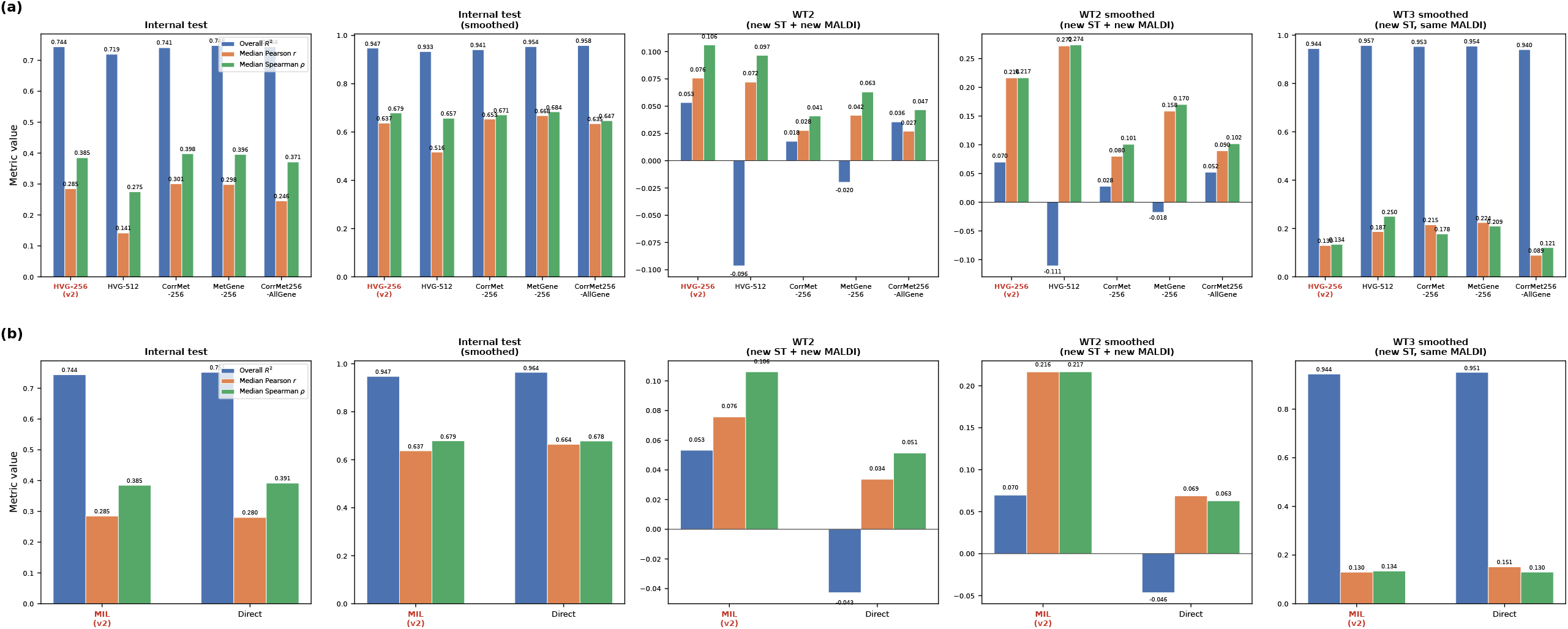
Systematic ablation identifies HVG-256 with MIL supervision as the only strategy that generalizes across sections. Performance of CHIMERA variants on three cohorts of increasing distributional shift. Each cohort is scored by overall spot-level R^2^ (blue), per-metabolite median Pearson r (orange) and median Spearman ρ (green), on raw and predictions. A, Five gene-feature strategies sharing the same backbone: HVG-256 (primary model, red label), HVG-512, CorrMet-256, MetGene-256 and CorrMet256-AllGene. HVG-256 is the only strategy retaining positive R^2^ on WT2. B, MIL versus Direct supervision at the HVG-256 backbone. Direct fits the internal fold marginally better but collapses on WT2, whereas MIL retains positive R^2^ across both held-out cohorts. Values are printed above each bar; y-axis ranges differ across panels.

On the internal fold, all strategies performed within a narrow band (*R*^2^ = 0.72– 0.75; median *r* = 0.25–0.30; median *ρ* = 0.28–0.40), converging further after smoothing (*R*^2^ ≥ 0.93; *ρ* ≥ 0.65; Fig. 2a, left panels). Within-sample evaluation was therefore non-discriminative, and differences among strategies emerged only under section shift.

The WT2 cohort—with both modalities acquired on new sections—separated the strategies sharply (Fig. 2a, middle panels). HVG-256 was the only route to retain simultaneously a positive overall *R*^2^ (0.053 raw; 0.070 smoothed) and positive median Pearson and Spearman correlations on both raw and smoothed predictions. HVG-512 inverted in sign (*R*^2^ = −0.096 raw; −0.111 smoothed) despite the highest smoothed Pearson (*r* = 0.272), indicating that a larger gene budget overfits magnitude while preserving rank order—consistent with a bias– variance collapse at 512 genes against a 189-dimensional metabolic output. The three metabolism-informed variants (CorrMet-256, MetGene-256, CorrMet256-AllGene) yielded negative *R*^2^ or Pearson *r* ≤ 0.08, showing that knowledge-based gene priors fail to transfer across sections. On WT3 (new Visium, shared MALDI-MSI with WT1) the ordering partially reversed: HVG-512 led on smoothed Pearson (*r* = 0.187) and Spearman (*ρ* = 0.250), while HVG-256 remained competitive (0.130; 0.134; Fig. 2a, right panel). HVG-256 is thus the unique strategy that preserves positive *R*^2^ on both held-out cohorts.

Holding the backbone at HVG-256, we next isolated the contribution of MIL supervision by comparing it with a Direct variant in which the pseudo-cell encoder was trained directly against the observed spot metabolome, bypassing the Π-weighted mixing step (Fig. 2b). Direct narrowly outperformed MIL on the internal fold (raw *R*^2^ 0.751 vs 0.744; smoothed 0.964 vs 0.947), but the ordering inverted sharply under section shift. On WT2, Direct collapsed to *R*^2^ = −0.043 while MIL held *R*^2^ = +0.053; after smoothing, MIL led by 0.117 in *R*^2^ (+0.070 vs −0.046), tripled the median Pearson (0.216 vs 0.069) and more than tripled the Spearman (0.217 vs 0.063). On WT3-smoothed the two were essentially tied (*R*^2^ 0.944 vs 0.951). Direct supervision thus fits the training distribution more aggressively but fails to transfer, whereas the Π-weighted mixing acts as a regularizer that prevents the encoder from exploiting section-specific covariance, at a modest within-sample fit penalty.

No single cohort resolves the design space on its own: the internal fold saturates for all strategies, WT2 alone favours HVG-256, and WT3 alone favours HVG-512. Only HVG-256 paired with MIL supervision remains competitive across all three cohorts simultaneously (Fig. 2a,b, red labels), indicating that cross-section transfer requires both a compact, knowledge-agnostic gene tokenization and a mixing-based regularizer. We therefore adopt HVG-256 + MIL as the primary CHIMERA model for all downstream analyses.

### CHIMERA predicts spot-level metabolomes and preserves spatial organization across sections

Under the primary HVG-256 + MIL configuration, the model converged within ∼100 epochs of training with no evidence of overfitting (train and validation loss trajectories remained parallel; Fig. 3A, B). On the combined WT+KO test fold (7,789 spots), the model attained an overall spot-level R^2^ of 0.752, with 123 of 189 metabolites (65%) showing R^2^ > 0. we also evaluated Pearson correlation on the identical fold. The per-spot correlations were uniformly high (median r = 0.882, n= 7,789 spots), indicating that the model faithfully reconstructs the relative metabolomic profile of each spot. At the per-metabolite level, 188 of 189 metabolites (99.5%) were positively correlated and 171 (90.5%) exceeded r = 0.1, with a median r = 0.285 (IQR 0.166_–_0.384) and 184 of 189 remaining significant (Fig 3C,table S2). the Ridge+MIL baseline underperformed CHIMERA on the test fold (per-metabolite median Pearson r = 0.177 vs 0.285). CHIMERA showed better Pearson r value in 158/189 metabolites, while baseline method performed better in 31/189 metabolites (Fig3 D,E). The model captures spot-level metabolomic structure with high fidelity.

**Figure 3:**
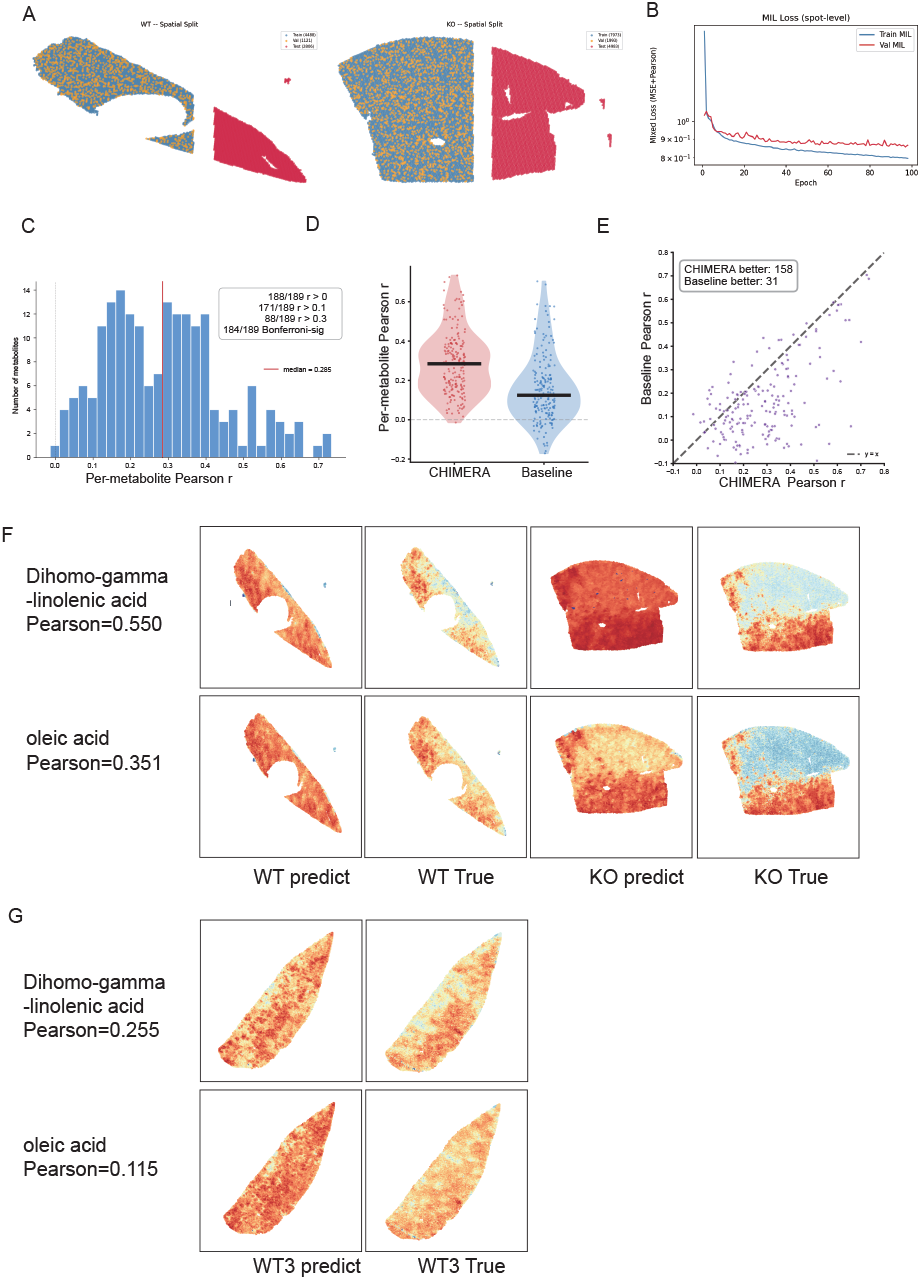
CHIMERA predicts spot-level metabolomes and preserves spatial organization across sections. A: Spatial cross-validation split of the WT (left) and KO (right) training sections into train, validation and test spots, with buffer gap zones (shown at the right of each section). B: Training (blue) and validation (red) MIL loss over training epochs. C: Distribution of per-metabolite Pearson r across the 189 metabolites on the combined WT+KO test fold (median r = 0.285; 188/189 positively correlated, 171/189 with r > 0.1, 184/189 significant after Bonferroni correction). D,E : Violin plot compare Pearson r between CHIMERA and baseline. F: Spatial distributions of predicted and measured dihomo-γ-linolenic acid (top; WT r = 0.550) and oleic acid (bottom; WT r = 0.351) on the WT and KO training sections. G: The same two metabolites on the WT3 section (DGLA r = 0.255; oleic acid r = 0.115)

To test whether the learned mapping reproduces spatially organized metabolite distributions rather than merely average abundances, we applied the trained encoder to the scRNA-seq reference and projected the predicted metabolomes back onto the corresponding Visium coordinates for side-by-side comparison with paired MALDI-MSI. We focused on dihomo-γ-linolenic acid (DGLA) and oleic acid, two lipids central to hepatic polyunsaturated and monounsaturated fatty acid metabolism and among the highest-confidence predictions on the training cohort (Pearson *r* = 0.52 and 0.48, respectively). On both WT and KO sections, the predicted spatial distributions of DGLA and oleic acid closely tracked the measured patterns (Fig. 3F)

We next asked whether this spatial fidelity survives the most stringent transfer setting, the held-out WT3 section in which both the Visium and the prediction are entirely unseen during training. Although spot-level Pearson correlations were attenuated as expected under distributional shift (DGLA *r* = 0.255; oleic acid *r* = 0.115), the dominant spatial organization of oleic acid and DGLA—was qualitatively preserved (Fig. 3G).

Together, these results demonstrate that CHIMERA delivers both quantitatively accurate spot-level predictions within sample and qualitatively faithful spatial reconstructions across sections, establishing the basis for single-cell-level metabolome inference on unpaired scRNA-seq data.

### CHIMERA recapitulates NAFLD-associated metabolic remodelling under Western diet

we next asked whether the learned transcriptome–metabolome mapping transfers to scRNA-seq data acquired independently of the training cohort.We applied CHIMERA to a Liver Cell Atlas cohort of 33241 CD45^-^ hepatic cells profiled under standard and Western diets (NDWD)[30], testing whether CHIMERA recovers chronic dietary metabolic remodelling.

Transcriptomic UMAP of the integrated atlas resolved five canonical hepatic cell types—hepatocytes, cholangiocytes, endothelial cells, hepatic stellate cells (HSCs) and hepatic stem/progenitor cells—with only minor shifts in global structure between ND and WD (Fig. 4a). UMAP computed on the CHIMERA-predicted metabolomes of the same cells recovered a comparable cell-type organization while redistributing cell density between conditions (Fig. 4b), indicating that diet-driven variation is reflect-able on the metabolic layer.

**Figure 4:**
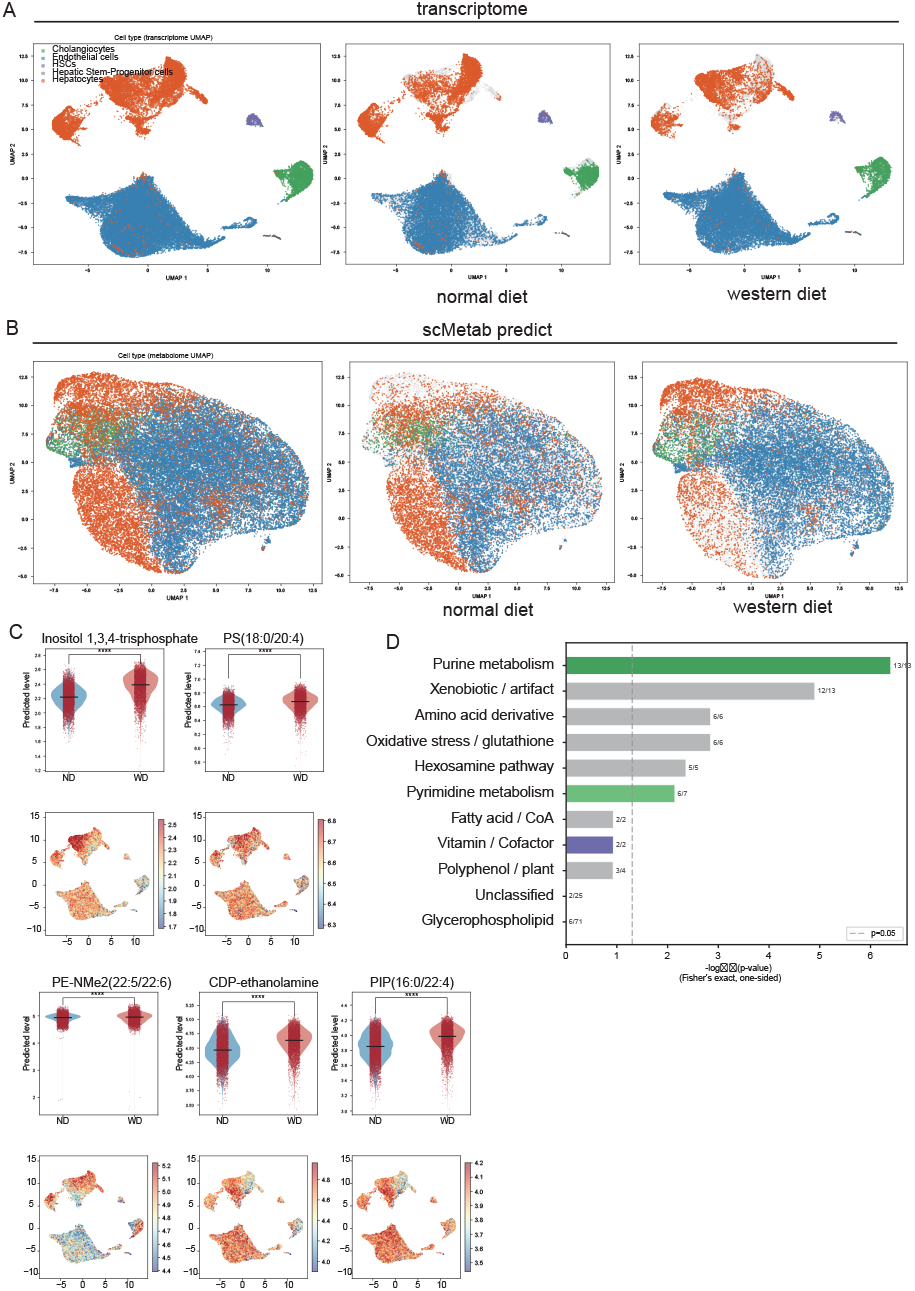
External validation on the Liver Cell Atlas normal-diet vs Western-diet cohort. CHIMERA was applied to 33241 hepatic cells spanning four conditions (SD/WD × 24w/36w). **A:** Transcriptomic UMAP coloured by cell type, shown for all cells (left), ND only (middle) and WD only (right). **B:** UMAP of CHIMERA-predicted metabolomes for the same cells, with matched panels. **C:** Predicted abundance of the five most reproducibly WD-up-regulated hepatocyte metabolites (violin plots, ND vs WD; UMAP feature plots below): inositol-1,3,4-trisphosphate, PS(18:0/20:4), PE-NMe2(22:5/22:6), CDP-ethanolamine and PIP(16:0/22:4). \*\*\*\**P* < 10^-^□, Wilcoxon rank-sum with Benjamini–Hochberg correction. **D:** Functional-category enrichment of WD-up metabolites (Fisher’s exact, one-sided; dashed line, *P* = 0.05). Fractions next to bars indicate hits / category size.

Differential abundance testing between WD and SD hepatocytes (Wilcoxon rank-sum with Benjamini–Hochberg correction; FDR < 0.05, |log□FC| > 0.1) identified top five metabolites with the largest and most reproducible WD-up signals across both 24- and 36-week cohorts: inositol-1,3,4-trisphosphate (a PI-signalling intermediate), phosphatidylserine PS(18:0/20:4), phosphatidylinositol-phosphate PIP(16:0/22:4), phosphatidylethanolamine-N,N-dimethyl PE-NMe2(22:5/22:6), and CDP-ethanolamine (a Kennedy-pathway precursor) (Fig. 4c,Table S3). The four phospholipid species are central intermediates of the glycerophospholipid-remodelling programme previously reported as a hallmark of human and murine NAFLD lipidomics, with hepatic and circulating PS, PI and PE all showing stage-dependent accumulation[31]. All five effect sizes were larger at 36 weeks than at 24 weeks, consistent with progressive diet-induced lipid remodelling characteristic of NAFLD progression[32].

Functional-category enrichment over the WD-up hepatocyte metabolite set revealed significant over-representation of purine metabolism (P = 3.97 × 10^-7^), oxidative-stress-associated species (P = 1.4 × 10^-3^), hexosamine biosynthesis and pyrimidine metabolism (Fig. 4d). The purine-metabolism signature is particularly notable: it recapitulates the established uric-acid/xanthine axis that drives NAFLD progression under hyper-caloric diet, validating model-derived predictions against a mechanistically established pathway[33-35]. The hexosamine biosynthesis enrichment is consistent with reports that hepatic HBP flux and protein O-GlcNAcylation are upregulated in NAFLD and promote de novo lipogenesis, ER stress and disease progression[36].

Together, these results demonstrate that Chimera is able to reconstructs metabolomic profiles from an entirely unrelated scRNA-seq dataset while preserving both within- and between-cell-type metabolic variation. Chimera recovered diet-associated metabolic alterations align with established NAFLD pathophysiology, further supporting its validity.

### Chimera reveals an Rarres2-induced metabolic alteration module in tumor-associated macrophages

To test whether single-cell metabolome prediction can uncover cell-state-resolved phenotypes inaccessible to single-modality analysis, we applied Chimera to 40,995 cells from a mouse Rarres2 (chemerin) knock-down hepatocellular carcinoma model. Rarres2 encodes chemerin, which binds CMKLR1 to recruit macrophages and natural killer cells; its role in hepatic tumor-associated macrophage (TAM) polarization and immune-metabolic coupling remains incompletely understood.

Among 8505 hepatic macrophages (annotated as Cd5l+, Vcam1+ or Spp1+ subtypes), the transcriptomic UMAP showed only modest separation between KD and NC cells (Fig. 5a,Fig S2), prompting us to ask whether Chimera-predicted metabolomes contained complementary information. Clustering macrophages on the top KD-vs-NC differentially abundant metabolites recovered four metabolic subclusters (MC-0 to MC-3; Fig. 5b) whose proportions differed markedly between conditions (χ^2^ = 88.2, P = 5.3 × 10^-19^; Fig. 5c). MC-3 is marked with cAMP, S-Formylglutathion, Adenosine thiamine diphosphate for metabolite and Selenbp1, Cd302, Sult1a1 for genes. MC-0/2 is marked with PE-NMe2, PS,PI for metabolite and Spp1, Mcm5, Tk1 for genes(Fig S2). Pathway annotation of cluster-marker genes and metabolites assigned each subcluster a distinct functional identity: MC-3 (n = 745) was dominated by lysosomal degradation and ether-lipid metabolism, the canonical signature of foamy lipid-associated macrophages (LAMs). MC-1 (n = 2,478) returned only six significant pathways with no active metabolic programme, consistent with a quiescent intermediate state. MC-0 (n = 3,395) was led by oxidative phosphorylation, the TCA cycle and proteasome activity across 50 pathways — a high-energy producing state. MC-2 (n = 1,887) was the most broadly activated, with 130 significant pathways spanning OXPHOS, proteasome, ribosome and ER protein processing(Figure S3, Table S4); combined with its Spp1+ enrichment, MC-2 corresponds to the classical SPP1+ like TAM end-state of hepatocellular carcinoma [37].

**Figure 5:**
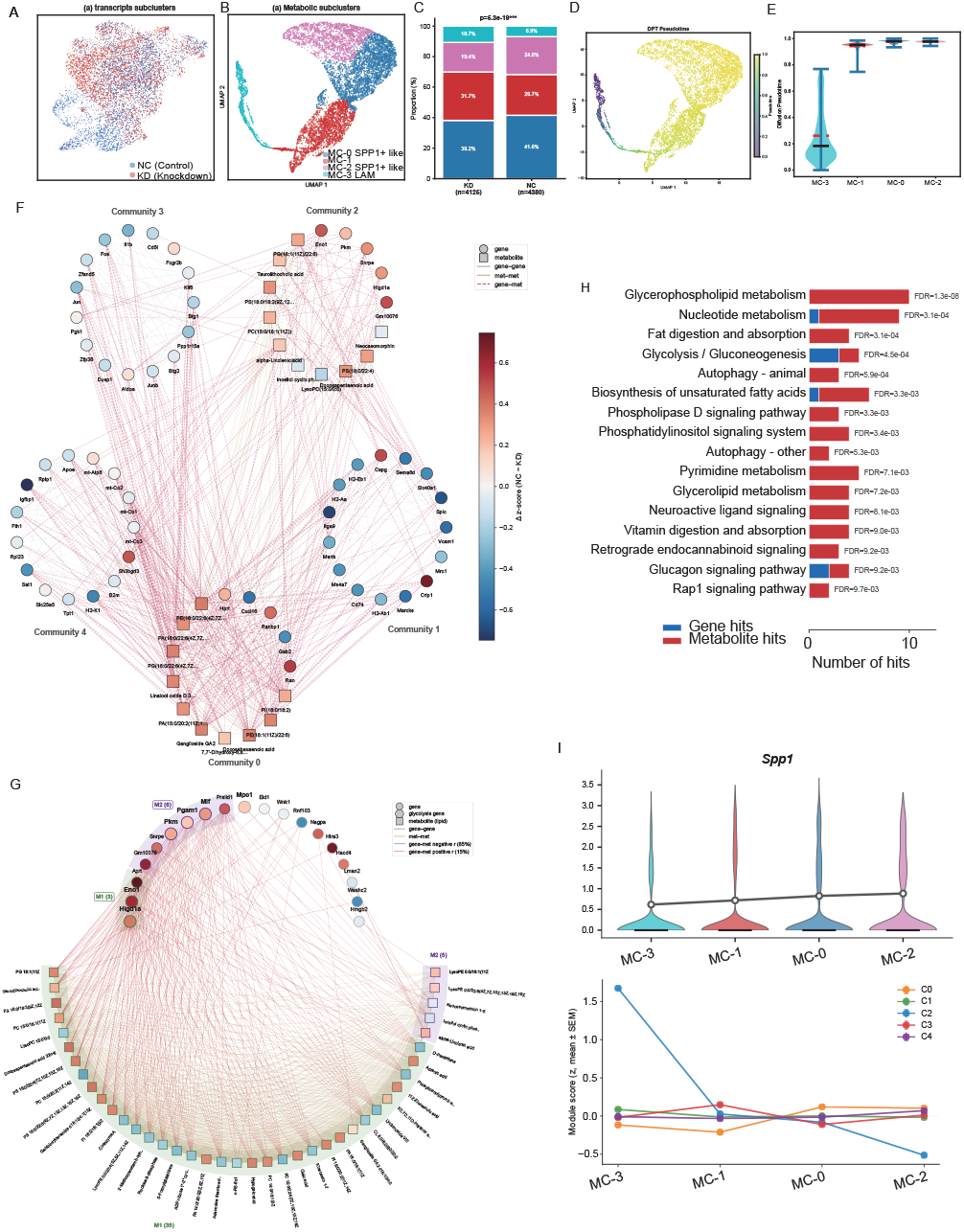
Chimera reveals an Rarres2-induced metabolic alteration module in tumor-associated macrophages. A: Transcriptomic UMAP of 8,505 hepatic macrophages coloured by condition (NC, n = 4,380; KD, n = 4,125). B: UMAP computed on CHIMERA-predicted metabolomes of the same cells, coloured by metabolic subcluster (MC-0 to MC-3). C: Stacked bar chart of metabolic-subcluster proportions by condition (χ^2^ = 88.2, P = 5.3 × 10^-19^). D: Diffusion pseudotime projected onto the metabolome UMAP, rooted at MC-3. E: Pseudotime distribution per subcluster. F: Heterogeneous gene–metabolite co-expression network partitioned by community detection into five communities (C0–C4); nodes are genes (circles) and metabolites (squares), coloured by KD-vs-NC effect size. G: MCODE refinement of the cross-modal C2 community, resolving two tightly connected sub-clusters (M1, the broad lipid pool; M2, the glycolytic core). H: Joint KEGG pathway enrichment of the C2 module combining gene (blue) and metabolite (red) hits. I: Spp1 and C2 module score along the MC-3 → MC-1 → MC-0 → MC-2 trajectory, shown against communities C0, C1, C3 and C4 (mean ± s.e.m.). *P < 0.05, **P < 0.01, ***P < 0.001.

Rarres2 expression shifted the metabolic polarization of TAMs from MC-3 toward MC-0/MC-2. Under KD, the MC-1 transition (26.7% → 31.7%) and MC-3 lipid-remodelling (6.9% → 10.7%) clusters expanded, while MC-0 (41.6% → 38.2%) and MC-2 (24.8% → 19.4%) contracted. Diffusion pseudotime organized the four subclusters into a polarization trajectory MC-3 → MC-1 → MC-0 → MC-2 (Fig. 5d, e), with KD cells concentrated at the MC-3 and NC cells progressing toward the MC-2 end-state. Along this axis, transcriptomic and metabolomic evidence converged on a single coordinated reprogramming: glycolysis (Eno1, Gapdh, Aldoa, Pkm), OXPHOS (Cox7b, Uqcrc1, Atp5c1) and de novo lipogenesis (Acly, Scd1, Fasn) transcripts rose monotonically; long-chain fatty acids accumulated in parallel; and NAD+/cofactor, purine and pyrimidine pools were progressively drawn down(Fig. S4). Together, these results support a model in which chemerin/CMKLR1 signalling drives TAMs from MC-3 toward MC-0/MC-2, coupling polarization to coordinated lipogenic, glycolytic and oxidative activation.

Chimera further enables co-embedding of the transcriptome and metabolome to identify co-regulated gene-metabolite modules. Building a heterogeneous co-expression network of gene-gene, metabolite-metabolite and cross-layer edges, and partitioning it with community detection, we recovered five gene-metabolite communities (C0–C4; Fig. 5f). C0 (652 genes + 43 metabolites) was dominated by translation, mitochondrial respiration and DHA-phospholipids; C1 (240 genes, 0 metabolites) carried the iron-recycling/MHC-II axis (Spic, Slc40a1, Vcam1); C3 (115 genes) and C4 (91 genes) encoded AP-1/NF-κB inflammation and ribosomal biosynthesis, respectively (Fig. 5f, Table S5).

Among these, C2 stood out as the only true cross-modal “Chimera” module, comprising 19 genes and 146 metabolites — a topology indicating that the gene and metabolite are potentially co-regulated. MCODE refinement of C2 resolved two tightly-connected sub-clusters (Fig. 5g): M1 (38 nodes), harbouring the metabolic genes Higd1a, Eno1 and Aprt together with 35 PUFA-phospholipids, gangliosides and free fatty acids — the broad lipid pool; and M2 (11 nodes), centred on the glycolytic core Pkm/Pgam1/Mif together with five lipid-degradation signalling species (LysoPE, α-linolenic acid and inositol cyclic phosphate). Joint KEGG-pathway analysis combining both layers identified glycerophospholipid metabolism, nucleotide metabolism, glycolysis/gluconeogenesis, biosynthesis of unsaturated fatty acids and taurine/hypotaurine metabolism as the principal hits (Fig. 5h).

The C2 module score was significantly higher in NC than in KD macrophages, confirming that Rarres2 expression maintains this metabolic state. Strikingly, the C2 score followed a strictly monotonic decrease along the MC-3 → MC-1 → MC-0 → MC-2 polarization trajectory (z: +1.67 → +0.03 → −0.08 → −0.51; FDR = 8.3 × 10^-163^), while Spp1 followed a monotonic increase(Fig. 5i). in marked contrast to communities C0/C1/C3/C4 which remained near baseline.

Taken together, these findings establish Rarres2/chemerin signalling reprogramming hepatic macrophages from a quiescent LAM-like state into a lipogenic, glycolytic, pro-inflammatory Spp1+ like phenotype, probably by altering the C2 gene-metabolite module. demonstrating that Chimera enable joint embedding of metabolite and genes, resolve coupled metabolic-transcriptic modules invisible to either modality alone.

## Discussion

We present CHIMERA, a transfer-learning framework that generates single-cell metabolomes from scRNA-seq data by learning transcriptome-to-metabolome relationships from spatially co-registered multi-omics measurements. By pairing each cell’s predicted metabolome with its measured transcriptome, CHIMERA reconstructs a single-cell multi-omics view that enables the discovery of differential metabolites and the co-embedding of genes and metabolites— analyses previously inaccessible without joint single-cell profiling. We demonstrate this across three settings: an internal benchmark on paired murine liver sections, external validation on an independent NDWD cohort in which CHIMERA recovered canonical NAFLD metabolite signatures (phosphoinositides, phosphatidylserines, CDP-ethanolamine and purine metabolism), and a mechanistic application to a Rarres2-KD HCC model, where it uncovered a coordinated lipogenic–glycolytic programme in tumour-associated macrophages—a dual-omics phenotype undetectable by either modality alone. CHIMERA thus turns the expanding universe of transcriptome-only atlases into a substrate for single-cell metabolic analysis, opening joint transcriptomic– metabolomic investigation without additional experimental cost.

CHIMERA differs fundamentally from existing computational metabolic inference approaches such as Compass, scFEA, METAFlux, scMetabolism, ssGSEA and MEBOCOST in three critical respects. First, it outputs metabolite abundance rather than relative fluxes or pathway-activity scores. Second, it learns transcriptome–metabolome relationships directly from experimental data, without constraining predictions to reactions encoded in KEGG or Recon. Third, by pairing predicted single-cell metabolomes with the measured transcriptome of the same cells, CHIMERA uniquely enables joint co-embedding of genes and metabolites in a shared latent space—an analysis that none of these methods supports. Such gene–metabolite coupling is invisible to either modality alone. The Rarres2-KD macrophage analysis illustrates this directly: CHIMERA recovers a co-regulated metabolic–transcript module (C2)—comprising glycolytic and lipid-metabolism genes alongside PUFA-phospholipids and free fatty acids— whose activity declines monotonically along a single macrophage polarization trajectory, a mechanistic coupling that knowledge-based tools, which score predefined pathways one at a time, cannot represent.

The external biological applications provide complementary lines of evidence that CHIMERA recovers genuine biochemical signal rather than training-cohort artefacts. In the NDWD cohort, the top predicted WD-up hepatocyte metabolites—phosphatidylserine, phosphatidylinositol-phosphate, CDP-ethanolamine and PE-NMe2—are central phospholipid-metabolism species whose accumulation has been documented in NAFLD lipidomic studies, and the concurrent purine-metabolism enrichment recapitulates the established uric-acid/xanthine axis of diet-induced steatosis. These signals strengthened monotonically from 24 to 36 weeks, mirroring the progressive dynamics of NAFLD pathogenesis. It is indicative that the inferred single-cell metabolome carries biologically faithful signal and supports downstream metabolic analysis.

Several limitations should be acknowledged. The pure-spectrum estimation step assumes that spot metabolomes decompose linearly into cell-type components weighted by deconvolution-derived proportions; this ignores paracrine metabolic exchange that may introduce non-linear contamination. Each Visium spot contains multiple cells, and deconvolution errors propagate into pure-spectrum and virtual-pure-spot constructions. MALDI-MSI annotation coverage is structurally biased towards lipid species (94 of 189 annotated metabolites in this study) and dependent on spectral databases, so predictions necessarily reflect a biased subset of the cellular metabolome; water-soluble glycolytic and TCA intermediates are particularly under-represented. Finally, the current model is trained and validated entirely on mouse liver; cross-tissue and cross-species transfer likely requires either multi-tissue training data or principled domain adaptation, and remains to be established.

In summary, CHIMERA is, to our knowledge, the first data-driven framework that outputs quantitative, database-independent single-cell metabolome predictions from scRNA-seq alone. Internal benchmarking, external validation in an independent NAFLD cohort, and mechanistic application to a *Rarres2*-KD HCC model collectively establish that CHIMERA recovers biologically interpretable metabolic phenotypes inaccessible to either experimental single-cell metabolomics or existing knowledge-based inference tools. We anticipate that applying CHIMERA across the expanding universe of scRNA-seq atlases will open a new dimension for single-cell biology—one in which every published transcriptomic dataset can be queried for its underlying metabolic logic.

## Declarations

### Ethics approval and consent to participate

The present study was performed in accordance with the Declaration of Helsinki. Approval for the use of human subjects was obtained from the research ethics committee of Huashan Hospital, Fudan University(KY2023-594), and informed consent was obtained from each individual enrolled in this study. All animal experiments were approved by the Animal Ethics Committee of Fudan University

### Consent for publication

Not applicable

### Availability of data and materials

The datasets used and/or analysed during the current study are available from the corresponding author on reasonable request.

## Acknowledgements

We sincerely thank Dr Chen-De Yang and Dr Dan-Ye in assisting establishing mouse model, Dr. Yang Zhang in assisting coding and writing improvement.

## Code and data availability

Trained model weights, preprocessing scripts and analysis notebooks are available at the project repository. The single-cell metabolome atlas (87,554 cells ×189 metabolites) is provided as AnnData and Seurat RDS files.

## Supplementary meterial

**Fig. S1 Overview of CHIMERA training data and cell-type annotation of the scRNA-seq reference**.

(A) Spatial multi-omics features of the three murine liver sections used for training and validation of CHIMERA (WT, WT2 and *Padi4*-knockout, KO).(B) UMAP of the murine liver scRNA-seq reference (n = 87,554 cells) used as the cellular basis for cell2location deconvolution.

**Fig. S2 Transcriptomic structure of KD and NC hepatic macrophages and marker metabolites/genes of the four metabolic subclusters (MC-0 – MC-3)**.

(A)Transcriptomic UMAPs of 8,505 hepatic macrophages from the *Rarres2* KD HCC cohort, shown separately for NC (control, left) and KD (*Rarres2* knock-down, middle)(B) Violin plots of single-cell expression (log1p CP10k) of the three subtype-defining genes *Cd5l, Vcam1* and *Spp1* across the three macrophage subtypes (n = 8,505 cells)(C) Marker-metabolite violin plots for the four metabolic subclusters (MC-0 – MC-3) (D) Marker-gene violin plots for the same four MCs.

**Fig. S3 Pathway activity per metabolic subcluster of hepatic macrophages**.

Heatmap of pathway activity across the four CHIMERA-derived macrophage metabolic subclusters (MC-0 – MC-3) in the *Rarres2* KD HCC cohort (n = 8,505 macrophages). asterisks denote two-sided Wilcoxon rank-sum significance with Benjamini–Hochberg correction (*FDR < 0.05; **FDR < 0.01; ***FDR < 0.001).

**Fig. S4 Coordinated metabolite and transcript reprogramming along the KD** → **NC polarization pseudotime in hepatic macrophages**.

(A)Metabolite-level changes along the trajectory, grouped into three biochemical categories. NAD□ & Cofactor metabolites (top), Purine & Pyrimidine metabolites (middle), Fatty Acid metabolites (bottom) (B) Transcript-level changes along the same pseudotime axis, grouped into three pathway modules. Glycolysis transcripts (top), OXPHOS transcripts (middle), Fatty-acid synthesis transcripts (bottom) Asterisks (where shown next to bars) denote significance of the underlying Spearman test (*P < 0.05; **P < 0.01; ***P < 0.001).

Table S1: gene lists of HVG-256 (v2), HVG-512 (v2_512),CorrMet-256,MetGene-256 and CorrMet256-AllGene.

Table S2: Per-metabolite prediction performance of CHIMERA on the test fold

Table S3: Differential abundance of CHIMERA-predicted metabolites between Western diet and standard diet hepatocytes in the Liver Cell Atlas NDWD cohort.

Table S4: pathway annotation of marker genes and marker metabolites for each macrophage metabolic cluster.

Table S5: Gene–metabolite co-embedding modules and functional annotation of hepatic macrophages

